# Modelling vasopressin synthesis and storage dynamics during prolonged osmotic challenge and recovery based on activity dependent upregulation of mRNA transcription

**DOI:** 10.1101/2022.06.27.497653

**Authors:** Duncan J. MacGregor

## Abstract

Hypothalamic vasopressin neurons are neuroendocrine cells which form part of the homeostatic systems that maintain osmotic pressure. In response to synaptic inputs encoding osmotic pressure and changes in plasma volume, they generate spike triggered secretion of peptide hormone vasopressin from axonal terminals in the posterior pituitary. The thousands of neurons’ secretory signals generate a summed plasma vasopressin signal acting at the kidneys to regulate water loss. Vasopressin is synthesised in cell bodies, packaged into vesicles, and transported to large stores in the pituitary terminals. Supported by activity-dependent upregulation of synthesis and transport, these stores can maintain a secretion response for several days of elevated osmotic pressure, tested by dehydration or salt loading. However, despite upregulated synthesis, stores gradually decline during sustained challenge, followed by a slow recovery. With no evidence of a store encoding feedback signal, previous modelling explained these synthesis dynamics based on activity-dependent upregulation of transcription and mRNA content. Here this model is adapted and integrated into our existing spiking and secretion model to generate a neuronal population model, able to simulate the secretion, store depletion, and replenishment, response to sustained osmotic challenge, matching the dynamics observed experimentally and making functional predictions for the cell body mechanisms.

## Introduction

Magnocellular vasopressin neurons, of the supraoptic and paraventricular nuclei of the hypothalamus, in response to input signals that encode osmotic pressure and plasma volume, synthesise and secrete the antidiuretic hormone vasopressin. Vasopressin in its antidiuretic role is a core element of the homeostatic system that maintains osmotic pressure (water/salt balance), signalling the kidneys to regulate how much water is retained. Acting as a heterogeneous population, these neurons are able to maintain a constantly functioning physiological signal over lifelong periods of time. To sustain such a signal the system must be both very robust and efficient. It must also be able to respond rapidly to large changes in demand.

Vasopressin is synthesised in the neuronal cell bodies and packaged into large dense core vesicles that are transported down the axons to the posterior pituitary, where the vesicles are stored in axon swellings and terminals that form larger reserve and smaller releasable pools. We have previously modelled the spiking and secretion mechanisms of these neurons (MacGregor and Leng, 2012, 2013), including the dynamics of these pools. Here we build and integrate with our existing model, a quantitative model of the synthesis mechanisms, to better understand how the properties of these neurons relate to their function on long timescales.

In normal (basal) conditions mammals drink and ingest sodium (in the diet) intermittently, but constantly lose water through respiration and perspiration. Under homeostatic regulation, osmotic pressure fluctuates around a ‘set point’; increases above this will be corrected by increased sodium excretion in urine, by increased thirst, and to compensate lack of availability or intermittent ingestion of water, by secreting vasopressin to concentrate the urine and minimise water loss. Falling below this set point occurs less commonly, when excess water has been consumed, or salt has been lost. Both tonic signalling and a response to perturbations must be maintained, and accordingly there is an almost continuous depletion of the pituitary vasopressin stores, which must be replenished by the synthesis, packaging, and transport of new vasopressin vesicles.

In conscious, normally hydrated rats, as in humans, the basal vasopressin plasma concentration is ~1 pg/ml (Robertson, Shelton and Athar, 1976; Verbalis, Baldwin and Robinson, 1986) and maximal antidiuresis is observed at a concentration of about 10 pg/ml. At concentrations higher than this, vasopressin continues to have an important role by its vasoconstrictor actions, which compensate for loss of fluid volume in conditions of dehydration. The pituitary vasopressin stores are large, between 1 and 2 μg (Leng and Ludwig, 2008), sufficient to maintain basal levels for around a month (Jones and Pickering, 1972). These large stores buffer against rapid increases in demand, but the rate of synthesis is also activity dependent, upregulating production in response to sustained increase in demand, and downregulating production in response to sustained low demand (Verbalis, Baldwin and Robinson, 1986). The system thus attempts to match supply and demand, minimising waste whilst protecting its stores (MacGregor, Clayton and Leng, 2013).

Upregulation of synthesis is limited however, and under conditions of sustained high demand, such as limited water access, the stores are depleted, falling to less than 30% after five days (Jones and Pickering, 1969). The activity-dependent synthesis rate is no longer able to match activity dependent secretion. When demand and the stimulating osmotic signal has returned to normal, the stores are gradually replenished, over a course of days (Young and Van Dyke, 1968). At this point the rate of synthesis must be exceeding the activity-dependent secretion rate.

Building a system, the simplest way to do this would be to have some feedback signal of store depletion. However we have no evidence for such a signal. The pituitary stores at the neurons’ secretory terminals are distant from the cell body and highly distributed among thousands of release sites, making the measuring and transmission of such a signal very difficult. The alternative is that some property of the synthesis mechanisms forms a memory of the challenge, sufficient to maintain higher synthesis rates beyond the direct stimulus. The best candidate for this is the pool of vasopressin mRNA. The mRNA pool increases in size several fold in response to prolonged challenge (Sherman, McKelvy and Watson, 1986). This mechanism was extensively investigated using both experimental and modelling work by a group in Pittsburgh in the late 1980s and early 1990s (Robinson *et al*., 1989; Fitzsimmons *et al*., 1992; Robinson and Fitzsimmons, 1993). They tested several alternative models (Fitzsimmons *et al*., 1992) and showed that the best match to observed dynamics of store depletion and replenishment uses an mRNA pool dependent rate of synthesis, combined with activity dependent upregulation of transcription. During a prolonged challenge the simulated mRNA pool increases in size, and following, the pool is gradually depleted, sustaining increased synthesis sufficient to replenish the stores, without requiring any feedback signal.

Their model (Fitzsimmons *et al*., 1992) forms the basis for our work here. Focussed on testing different models of the mRNA pool and its relation to synthesis rate, they made the simplification that the synthesis rate always approaches a steady state equal to the rate of secretion. What limits this response and causes store depletion (and replenishment) in their model, is that this change in rate uses a long time constant, dependent on the half-life of mRNA, estimated by them at two days. They tested alternative models with activity dependent mRNA decay, and with longer and shorter fixed decay rates, but the best fit to the experimentally observed dynamics was with this model. The model was fitted to data from several sources (Young and Van Dyke, 1968; Jones and Pickering, 1969; Zingg, Lefebvre and Almazan, 1986; Roberts *et al*.,1991) measuring the changing vasopressin content during a prolonged osmotic challenge (water deprivation or salt loading through high Na^+^ drinking water), and the following recovery. It was also based on data estimating the rate of synthesis by measuring the accumulated vasopressin content at the cell bodies with transport blocked to the peripheral stores (Roberts *et al*., 1991). Synthesis rates were estimated to be ~1-3 ng/h at basal, and 10 ng/h under hyper-osmotic conditions (3 days of salt loading). They also showed that the synthesis rates only gradually return toward basal levels in the days following the osmotic challenge and that the transport rates (from cell body to peripheral stores) up- and down-regulate in parallel (Roberts *et al*., 1991). This prolonged upregulation of synthesis and transport acts to replenish the peripheral stores.

The Pittsburgh model, which only simulates synthesis and the store, using the simplification that synthesis always tracks secretion, does not deal with the pathway between osmotic stimulus and regulation of mRNA. A representation of this pathway is required for our model, which takes as input a synaptic signal that encodes osmotic pressure. It is well established that osmotic stimulation increases vasopressin mRNA content (Sherman, McKelvy and Watson, 1986), and also known that hypo-osmotic conditions reduce mRNA content (Svane *et al*., 1995). As well as increasing transcription rates, prolonged osmotic stimulation increases the length of the mRNA poly(A) tails (Carrazana, Pasieka and Majzoub, 1988; Zingg, Lefebvre and Almazan, 1988), and the overall changes in content are likely due to a combination. Longer poly(A) tails are thought to either increase mRNA stability or increase translation efficiency (Emanuel *et al*., 1998). The functional effect of either would be to increase the amount of synthesis per unit of mRNA.

The pathway between osmotic stimulus and transcription is still uncertain. The major candidate is a pathway via cyclic AMP (Carter and Murphy, 1989; Sladek *et al*., 1996; Wong *et al*., 2003) that acts to drive the CREB3L1 transcription promoter (Greenwood *et al*., 2015). There is also evidence for a glutamate-NMDA receptor-Ca^2+^ entry driven pathway (Lake, Corrêa and Müller, 2019). Here we are using a very simple representation to predict the necessary dynamics rather than any detailed modelling of the mechanisms.

Our objective here was to integrate, adapt and extend the Pittsburgh model into our existing integrated spiking and secretion model in order to fully simulate the pathway from osmotic signal to plasma hormone signal. The challenge we identified when testing the secretion model (MacGregor and Leng, 2013) is that heterogeneity, which brings essential benefits to producing a robust signal response, results in widely varying rates of secretion and store depletion across the population. The synthesis mechanism must be able to cope with varied demand not only as a population but also between individual neurons.

The new synthesis modelling has been kept as simple and general as possible, and should be capable of being adapted to other neuroendocrine cells, but is still able to produce strong quantitative as well as qualitative matches to the experimental data on synthesis rates, mRNA content, and depletion and repletion of vasopressin stores during prolonged osmotic challenge and recovery. However, in designing and fitting to match experimental data that shows a cycle of depletion and recovery during and after an osmotic challenge, the synthesis model is essentially constrained to fail at the task of matching supply to demand. By attempting to fix this in the model we explore why the stores get depleted; what are the limiting mechanisms, and why these limits might be necessary.

## Results

### Osmotic Challenge and Recovery Data

To set targets for fitting and testing the model, an extensive literature survey was used to gather multiple types of physiological data recorded in rats during a prolonged osmotic challenge and the following recovery. This extends the examples of (Fitzsimmons *et al*., 1992) where they compared multiple sources measuring the depletion and recovery of pituitary vasopressin stores. The data used here (Figure 1) includes osmolarity, plasma Na concentration, plasma vasopressin concentration, hypothalamic vasopressin mRNA content, and pituitary vasopressin content, during depletion and recovery. The data was extracted and reconstructed mostly by using graphics software (Adobe Illustrator) to measure points plotted in figures. A full list of the sources and tables of the data are given in the supplementary material.

**Figure 1.**
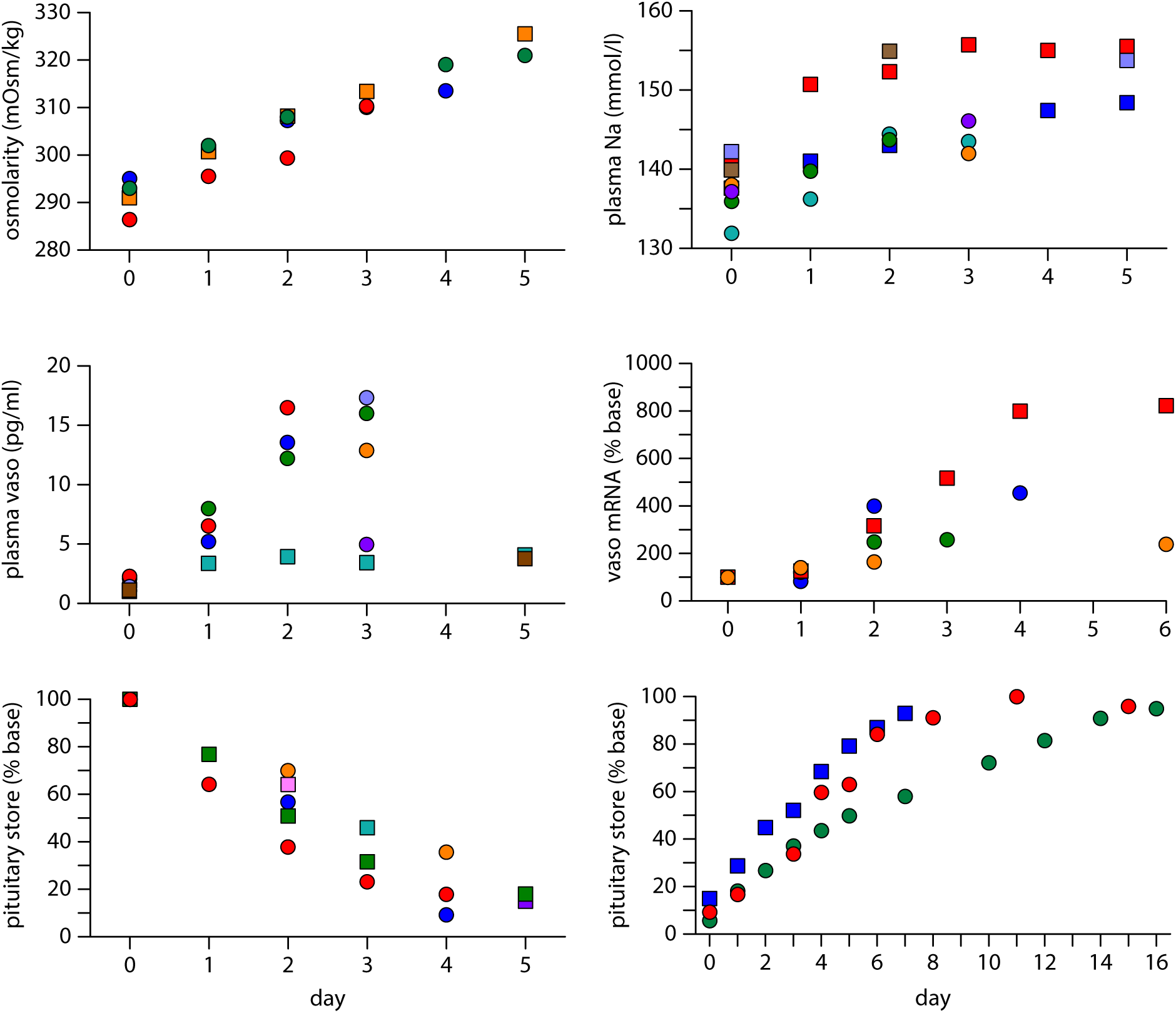
Experimental data gathered during prolonged osmotic challenge and recovery. The data here is gathered from multiple published sources where rats have been measured during a prolonged osmotic challenge consisting of several days of dehydration or salt loading (high sodium drinking water), and the following recovery period, with normal water access restored. During the challenge osmolarity and plasma Na (top panels) rise mostly linearly, with some data showing a reduced rate of rise and plateau towards day 4 and 5. Radioimmunoassay measured plasma vasopressin concentration (mid left) mostly shows a matching linear rise, but data is mostly limited to three days, and varies in magnitude between dehydration (higher) and salt loading (lower) protocols. Vasopressin cell body mRNA content (mid right) shows a mostly linear rise after one day that eventually plateaus. The data is variable, but the most consistent experiments, with more time points, suggest a five to eight fold rise in content. The core target data for the model is the measurements of pituitary store content (bottom). During the challenge there is a mostly linear fall in pituitary content, which slows after day 3, falling to around 15 to 30% of normal content. During recovery, where osmolarity rapidly (a few hours) returns to normal, the stores are slowly replenished over about two weeks. The faster recovery shown here (blue squares) is in rats made hypo-osmotic after the prolonged hyper-osmotic challenge. Detail on the sources is given in supplementary Figure S1.

The comparisons between multiple sources are imperfect. Experiments use different breeds and ages of rat, different timings of measurements (which are likely to have circadian sensitivity), and different assay techniques. In particular plasma Na is measured both by flame photometry and by electrode based techniques. Measurements of plasma vasopressin by radioimmunoassay are dependent on varying sample extraction techniques and assay antibodies. Using multiple sources has attempted to provide as clear a consensus as can be achieved, providing data to fit and test the input osmotic stimulus (osmolarity and plasma Na), the internal mRNA content, and the output plasma vasopressin and pituitary content.

The osmotic stimulus protocols vary between using dehydration (water deprivation) and salt loading (high Na drinking water) to raise osmolarity. In all the measurements except for plasma vasopressin these two protocols appear to produce an equivalent response (Figure 1). The lower plasma vasopressin concentrations observed under salt loading (~4 pg/ml vs 15 pg/ml under dehydration) are inconsistent with the similar rates of pituitary content depletion. Content depletion is likely to be a more robust measure of sustained vasopressin secretion rates, and so the model here targets the higher and more consistent plasma vasopressin concentrations observed under dehydration.

### The Spiking and Secretion Models

The spiking model used to generate the results here uses parameters (Table 1) chosen to simulate a typical magnocellular vasopressin neuron, based on detailed fits to *in vivo* recordings (MacGregor and Leng, 2012). As the synaptic input rate is increased, spiking shifts from silence to irregular spiking, phasic patterned spiking (long bursts and silences), increasing burst durations, and eventually continuous spiking. Figure 2 shows phasic spiking in the model. The non-linear stimulus-response properties of the secretory terminals (frequency facilitation and fatigue), simulated by the secretion model, are such that the phasic pattern is optimal in terms of secretion per spike. Thus the increase in the rate of secretion slows as the neuron is driven into less optimal continuous spiking.

**Table 1:** Spiking Model Parameters

| Name | Description | Value (units) |
| --- | --- | --- |
| $I_{re}$ | excitatory input rate | 230 (Hz) |
| $I_{ratio}$ | inhibitory input ratio | 0.75 |
| $e_h$ | EPSP amplitude | 3 (mV) |
| $i_h$ | IPSP amplitude | -3 (mV) |
| $\lambda_{syn}$ | PSP half life | 7.5 (ms) |
| $k_{HAP}$ | HAP amplitude per spike | 60 (mV) |
| $\lambda_{HAP}$ | HAP half life | 9 (ms) |
| $k_{DAP}$ | fast DAP amplitude per spike | 1 (mV) |
| $\lambda_{DAP}$ | fast DAP half life | 150 (ms) |
| $k_{AHP_{slow}}$ | slow AHP activation factor | 0.00012 (mV/nM) |
| $\lambda_{AHP_{slow}}$ | slow AHP half life | 10000 (ms) |
| $C_{AHP_{slow}}$ | minimum $[Ca]_i$ to activate slow AHP | 200 (nM) |
| $C_{rest}$ | rest $[Ca]_i$ | 113 (nM) |
| $k_C$ | $[Ca]_i$ increase per spike | 11 (nM) |
| $\lambda_C$ | $[Ca]_i$ half life | 2500 (ms) |
| $k_D$ | dynorphin activation per spike | 2.693 |
| $\lambda_D$ | dynorphin half life | 7500 (ms) |
| $k_L$ | $K^+$ leak calcium sensitivity | 36 (nM) |
| $g_L$ | $K^+$ leak maximum voltage | 8.5 (mV) |
| $V_{rest}$ | resting potential | -62 (mV) |
| $V_{thresh}$ | spike threshold potential | -50 (mV) |

**Figure 2.**
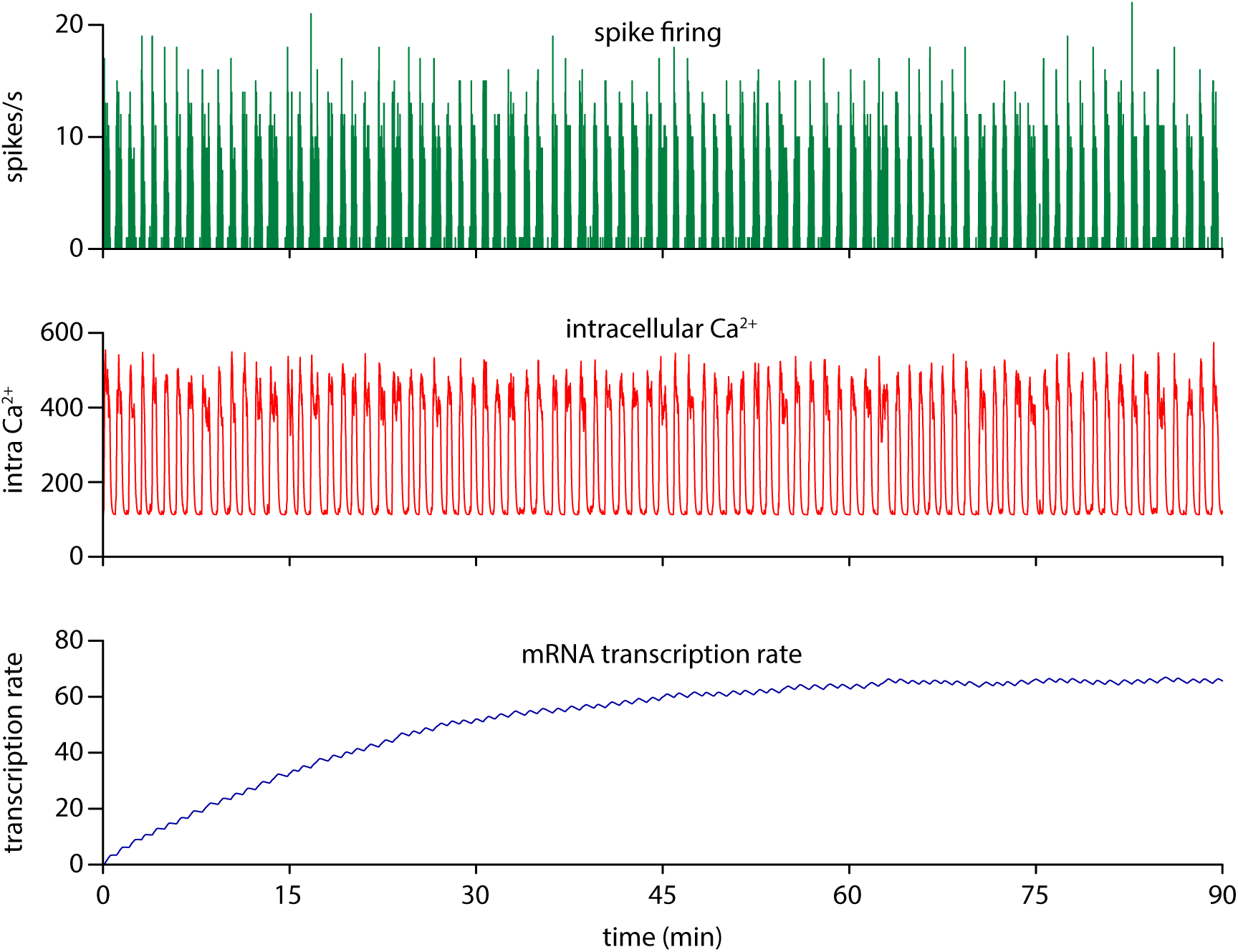
Spike activity dependent regulation of transcription. Phasic spiking in a highly stimulated integrate-and-fire based model neuron is both driven by and generates Ca^2+^ entry, producing an intracellular Ca^2+^ signal that is used to drive the model’s vasopressin mRNA transcription rate. The essential dynamic is that the mechanism translates the rapidly changing and noisy electrical activity into a sustained slow-changing measure of activity.

The secretion model is modified from the previously published version as described in the Methods. It is coupled to a model of plasma diffusion and clearance which we previously developed to simulate oxytocin plasma concentrations (Maícas-Royo, Leng and MacGregor, 2018), with parameters adjusted using experimental data on vasopressin plasma concentrations and clearance rates, again as described in the Methods.

### Synthesis Model Basic Function and Tuning

The synthesis model was initially tested using a single neuron. The secretion rate is scaled to the number of neurons to maintain comparable secretion and plasma concentrations independent of population size.

The transcription rate (*T*) half-life λ*_T_* = 1000 s and upregulation rate *k_T_* = 0.33 were chosen to produce a plausible timescale transcription rate signal, taking with input rate *I_re_* = 380 Hz, ~1 h to reach equilibrium *T* = ~ 60 (arbitrary units) from an initial *T* = 0 (Figure 2). This signal forms a long timescale measure of spike activity which in turn drives the increase in mRNA content.

Figure 3 illustrates the basic function of the model with steady input rate *I_re_* = 380 Hz, corresponding to a sustained hyperosmotic state. Vasopressin mRNA content rises very slowly towards an equilibrium at a rate and level determined by the balance between the transcription rate and depletion due to translation and synthesis. The rate of synthesis is directly proportional to the mRNA content (*m*). The secretion rate and plasma concentration driven by the single phasic neuron are noisy but sustain steady levels until the reserve store is depleted. The releasable pool (which is refilled from the reserve) buffers the secretion response to maintain a steady rate until the reserve store is very heavily depleted. With no synthesis, the store fully depletes and plasma concentration falls to zero. With synthesis, the rate is insufficient to match the highly stimulated secretion rate and the store is still depleted, though at a slower rate. When it is depleted, secretion and plasma concentration is sustained, at a level purely dependent on the upregulated synthesis rate. We would not expect to observe this in the heterogeneous population *in vivo*, but this is what we would predict in a homogeneous population, assuming a sustained osmotic stimulus.

**Figure 3.**
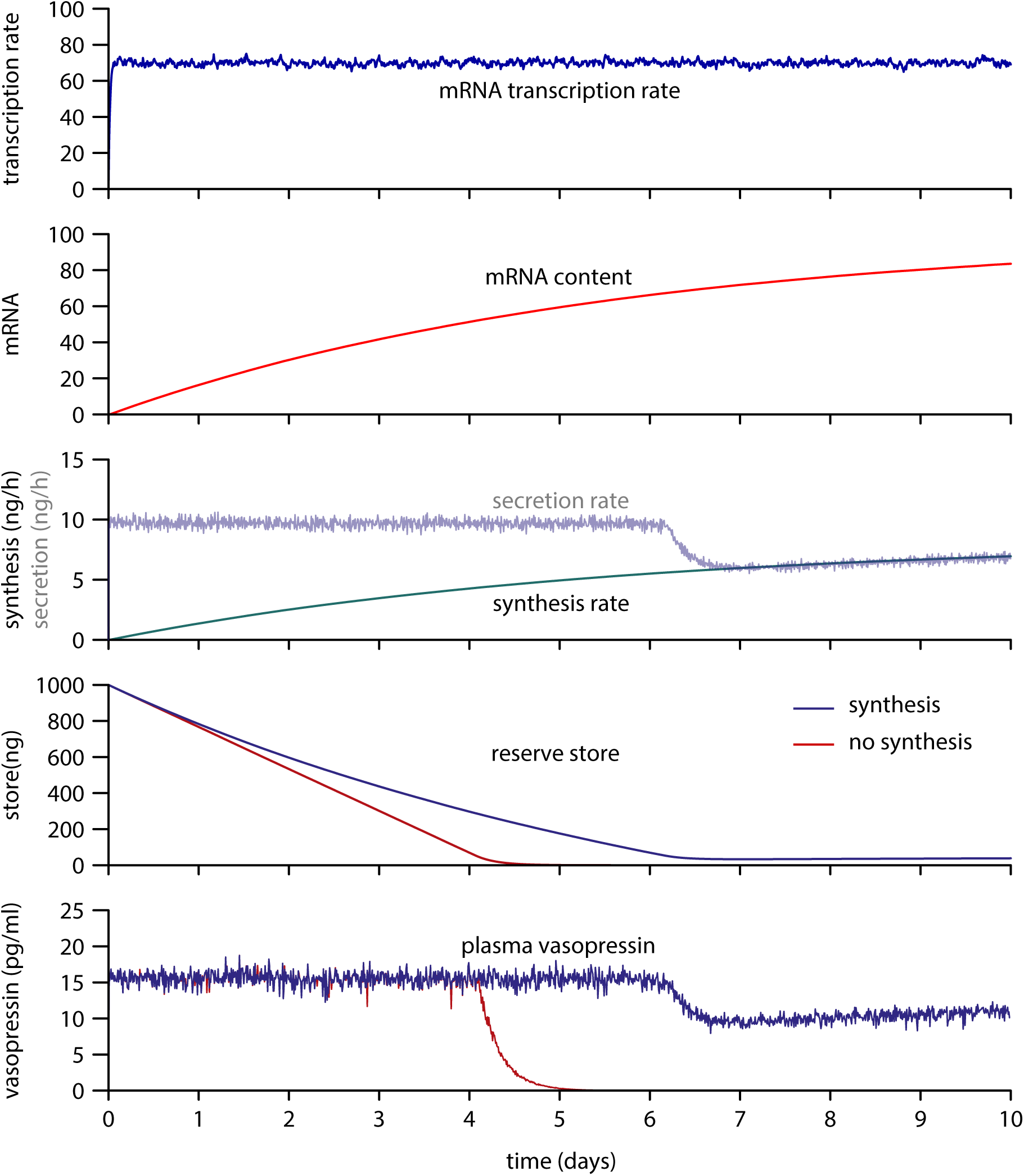
Single neuron transcription-dependent regulation of mRNA content and synthesis rates. For illustration, rather than physiological simulation, the model here is initialised with a full store and zero stimulus, switching at time 0 to a sustained highly osmotic input signal. Transcription drives the accumulation of mRNA content, which in turn determines the rate of synthesis which maintains (or slows the depletion of) hormone stores. In the rapid change to a highly stimulated state here, elevated synthesis is not sufficient to match the rate of secretion, and stores are gradually depleted. When the stores are depleted the rate of secretion becomes purely synthesis rate dependent.

The secretion model parameters were fixed by fitting the secretion model to *in vitro* data as described in the Methods and detailed in (MacGregor and Leng, 2013; Maícas-Royo, Leng and MacGregor, 2018). Coupled to the plasma model, this allows the prediction of the rates of secretion that correspond to plasma concentrations observed *in vivo*. In basal normo-osmotic conditions rat plasma vasopressin *in vivo* is ~ 1 pg/ml. In highly stimulated hyper-osmotic conditions plasma vasopressin *in vivo* rises to around 20 pg/ml. The left panels of Figure 4 show the single neuron model sustaining a mean 1 pg/ml plasma concentration with input rate *I_re_* = 252 Hz. The initial value for mRNA content (*m* = 15), was set using an initial test to find its stable value at this input rate. The first target for tuning the synthesis model parameters was for synthesis to match secretion in basal conditions, in order to sustain a stable reserve store. The reserve store plot shows this achieved by reducing the synthesis rate (*s_r_* = 0.65) compared to the final heterogeneous model parameter (*s_r_* = 1.1), which produces a small rise in the store, with synthesis exceeding secretion.

**Figure 4.**
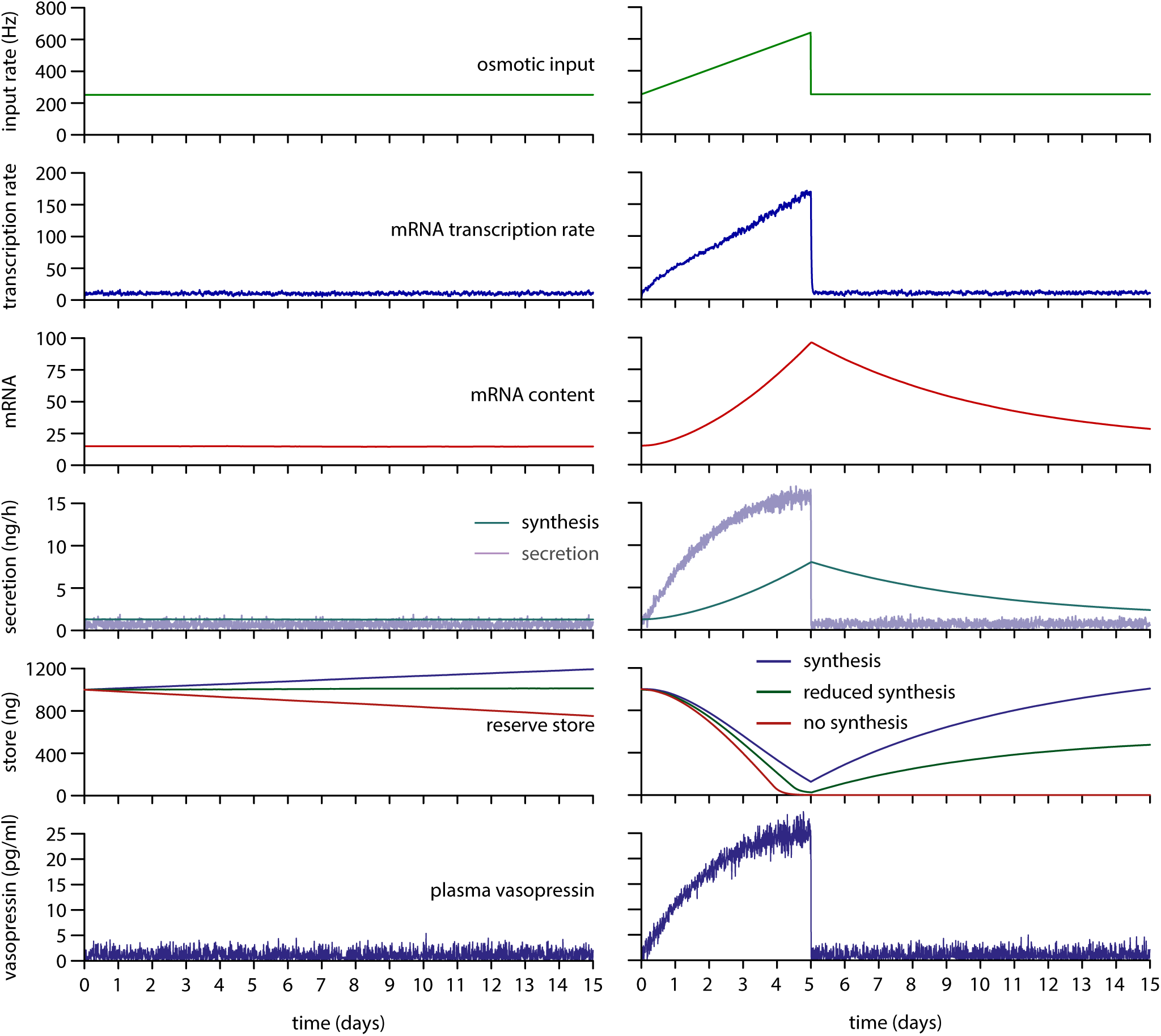
Single neuron basal activity and prolonged osmotic challenge and recovery. The plots on the left show basal activity sustaining a mean 1pg/ml plasma vasopressin. The transcription rate rapidly rises to sustain mRNA content at 15 units. With default synthesis rate scaling *s_r_* = 1.1, the synthesis rate slightly exceeds the secretion rate and the reserve store increases (blue). Setting *s_r_* = 0.65 (green) maintains a stable store. Removing synthesis by setting *s_r_* = 0 shows a depleting store. The plots on the right show a 5 day osmotic challenge (a linear ramp from basal, simulating progressive dehydration or salt loading) followed by 10 day recovery (input returned to basal). The transcription rate mostly tracks the osmotic stimulus with some drop off due to non-linearities in the spiking response. The mRNA content rises non-linearly as the balance shifts between transcription, and depletion due to synthesis (translation). The synthesis rate increases but fails to track the increasing secretion rate. The reserve store is gradually depleted to ~27%. Plasma vasopressin increases initially linearly but then slows as the neurons shift from phasic to continuous spiking. Following the ramped challenge, secretion falls to basal rates, but elevated synthesis is sustained by the increased mRNA content, depleting this to recharge the reserve store.

### Simulating Sustained Osmotic Challenge and Recovery

The second target for tuning the model was to match the experimental data measuring vasopressin store content in rats during a five day osmotic challenge (no water access, or salt loading using high Na^+^ drinking water) and the following recovery (Figure 1). Experiments measuring osmolarity (or equivalent plasma Na^+^) and plasma vasopressin during similar protocols (Walters and Hatton, 1974; Nordmann, 1985; Yue *et al*., 2008) suggest that the osmotic stimulus rises mostly linearly during the challenge for at least the first three days before levelling off at sustained high levels, and rapidly recovering after the challenge period. We simplified this by using a linear ramp in the input rate to simulate the prolonged osmotic challenge, illustrated in the right hand panels of Figure 4. The initial input rate 252 Hz was ramped to 640 Hz over 5 days and then returned to 252 Hz, targeted to match the store depletion observed in the experimental data.

The transcription rate mostly tracks the osmotic stimulus. The stores decline in content to ~ 25% before slowly recovering following the challenge, matching the experimental data and the results with the original Pittsburgh synthesis model. The mRNA content shows a more non-linear increase, and decline during the recovery period, as it sustains elevated synthesis rates to replenish the stores. However, the reduced synthesis rate (*s_r_* = 0.65) used to match secretion at basal levels (Figure 4 left) produces more depletion of the stores and an incomplete recovery. As shown below, the synthesis model was more difficult to tune for a homogeneous than a heterogeneous population.

### Synthesis Response in a Heterogeneous Population

Osmotically stimulated vasopressin neurons recorded *in vivo* show widely varying spiking rates. We can simulate this heterogeneity by randomly varying the amount of synaptic input received by each model neuron, applying a varied input density parameter, as detailed in the Methods. This heterogeneity has a substantial functional advantage, producing a much more linear plasma vasopressin response to a changing osmotic input signal than a homogeneous population (MacGregor and Leng, 2013). This matches the response that has been observed experimentally, however it results in also producing highly heterogeneous secretion rates and store depletion across the population. Here we tested how the synthesis model would respond to these varied stimulus and store depletion rates, and how this would affect the summed population response. The model was set up to record both the summed population and the individual neuron mRNA content, reserve store, and secretion rates.

A 100 neuron heterogeneous population was randomly generated with a lognormal distribution, illustrated in the inset of Figure 5. The basal population input rate (which is modified by each neuron’s input density) was set to 207 Hz to produce a sustained 1 pg/ml plasma vasopressin concentration. The initial values for *m* were set for each neuron by running the model with a 207 Hz population input until the neurons’*m* values had stabilised, starting with common values of *m* = 10. No other parameter adjustment was required from the same basal protocol tested with the homogeneous model (Figure 4). This produced both a stable plasma vasopressin signal and a stable summed reserve store. As well as the summed population data Figure 5 shows a sample of three neurons from the low, middle, and high end of the activity distribution. The low and middle neurons (blue and green) both had very low spiking and secretion rates. Their mRNA content *m* was down-regulated to almost zero, with a matching low synthesis rate. This matches what is observed experimentally in hypo-osmotic conditions, showing the ability to down-regulate as well as upregulate from the basal synthesis activity. The high activity neurons (example here in red) show an increased mRNA content and a sustained secretion rate much higher than the population mean. They also show a gradual increase of their stores as synthesis exceeds the secretion rate.

**Figure 5.**
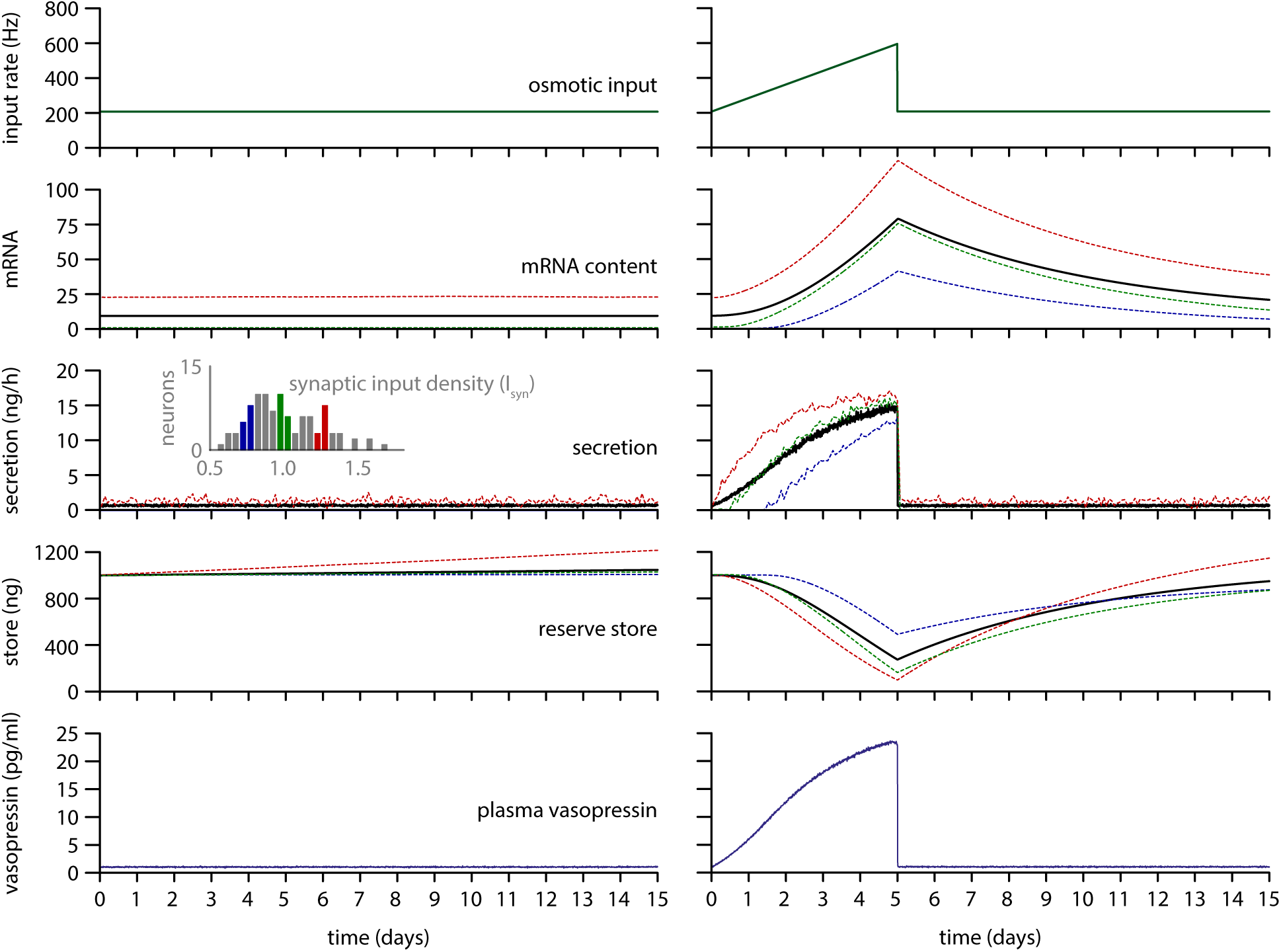
Basal activity and prolonged challenge and recovery in a heterogeneous population. The data here show a 100 neuron heterogeneous population following the same protocols as Figure 4. The inset distribution shows the heterogeneous input rates and the colour coded example neurons. In the left hand plots, with default synthesis scaling *s_r_* = 1.1, the heterogeneous population sustains a stable reserve store at basal activity (1 pg/ml plasma vasopressin). The majority of the secretion is from the more active neurons (red) and these show gradual increase of individual neuron stores. In the right hand plots, the challenge and recovery protocol shows similar results to the single neuron (Figure 4), but with a more stable and linear plasma signal. The individual neurons show some complex divergence in their store recovery, due to varied non-linearities in secretion and synthesis. The more active cells, even with highly elevated mRNA, show more rapid depletion, but also a more rapid, and even excess recovery.

Testing the ramped challenge and recovery protocol (initial population input rate 207 Hz ramped to 595 Hz over 5 days then returned to 207 Hz), plasma vasopressin and the summed population data shows very similar results to the homogeneous population. In this highly stimulated protocol the middle activity neuron more closely matches the mean population rates. The secretion rates of individual neurons are much more non-linear than the population mean, and the plasma concentration shows a more linear response to the ramped stimulus than the homogeneous population.

### Model Compared to Experimental Data

Figure 6 uses the same heterogeneous population model as Figure 5, with an extended time protocol for comparison with the experimental data, 2 days of basal activity followed by a 5 day osmotic challenge and 15 days of recovery. The results show a population store that falls to ~ 25 % during the 5 day challenge and replenishes to almost full over the 15 day recovery period, very similar to the experimental data in rats. A notable difference however is that the model’s store using the linear ramp (black) protocol shows a slower initial decline. The match was improved by using a more non-linear ramp (red) in the osmotic input signal, producing a more rapid initial increase that gradually slows. Experimental evidence for the ramp in osmotic input and plasma vasopressin is variable and limited by temporally sparse measurements but suggests something that lies between these two. The model was further modified (blue) by adding a 24 h delay between the synthesis rate and the store fill rate representing the slow transport of new vesicles from the cell body to the pituitary stores, thought to take up to 24 h depending on osmotic status (Russell, Brownstein and Gainer, 1981). The delay more closely matches the depletion observed experimentally, but its effect is modulatory, and not sufficient to explain store depletion alone.

**Figure 6.**
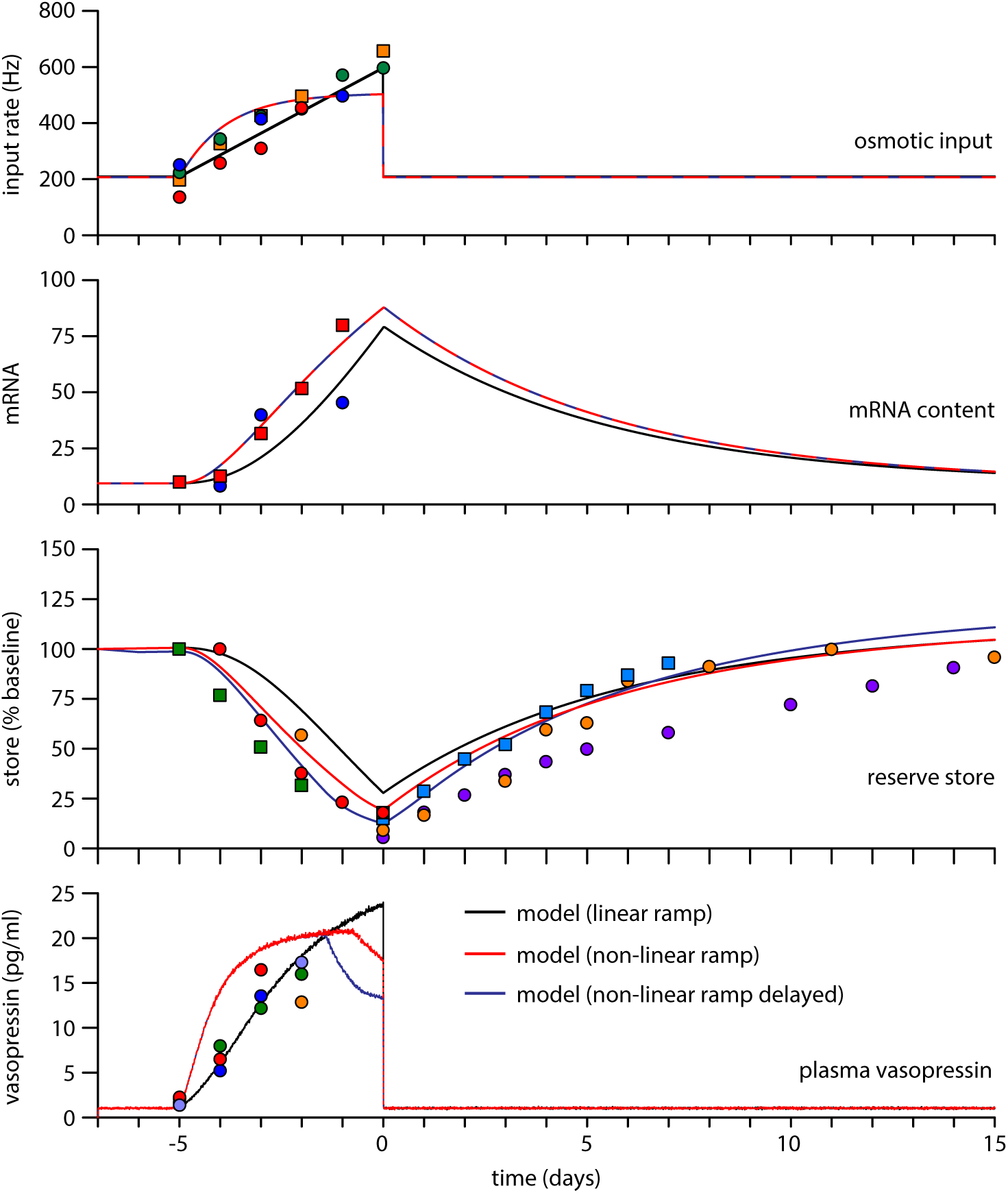
Model challenge and recovery compared to experimental data. The 100 neuron heterogeneous population here simulates 2 days of basal activity, followed by 5 days of dehydration, and 15 days of recovery, compared against experimental data from Figure 1. Two input signal protocols are compared, a default linear ramp (black) and a non-linear ramp (red) with a more rapid initial increase in osmotic signal. Within the variability of the experimental data both ramps are potentially consistent. The linear ramp varies from the store data in its slower initial decline in content, more closely matched by the non-linear ramp, which produces a faster increase in secretion than synthesis, resulting in more rapid initial store depletion. The model was further tested with an added delay between synthesis and store fill rate (blue), simulating the estimated 24h transport delay. The delay only moderately changes the population store depletion and recovery profile, however more active neurons become fully depleted, resulting in a drop off in the plasma signal.

### What Limits the Synthesis Response?

The current model matches the limited upregulation of synthesis, and depletion of stores, observed in the experimental data. Changes to the model, attempting to improve this response, were tested to predict which mechanisms might be responsible for this limited ability to match secretion demand (Figure 7). Two methods were found which were able to produce a much faster upregulation of synthesis while maintaining the ability to function in basal and stimulated states without under- or over-filling the stores. The first method (red in Figure 7) accelerates the upregulation of transcription. This required three parameter changes, increasing the rate of transcription but also compensating the increased amount so that only the speed of the response was changed (*k_T_* 0.33 to 3.3, *S_basai_* 0.7 to 7, *S_r_* 1.1 to 0.11). The produces a much faster increase in the mRNA store and corresponding synthesis rate, resulting in a much smaller vasopressin store depletion. However it also predicts a much larger increase in mRNA than is observed experimentally (20-fold compared to ~5 to 8-fold). The second method (blue in Figure 7) increases the rate of translation (*s_basai_* 0.7 to 3), increasing the rate of synthesis in exchange for a faster depletion of the mRNA store. This similarly produces a faster upregulation and reduced vasopressin store depletion but also results in a much smaller increase in the mRNA store, since the mRNA equilibrium level is determined by the balance between transcription and translation. Thus, the model predicts that the main elements responsible for vasopressin store depletion are the lag in upregulation of mRNA, and the maximum mRNA capacity, combined with a limit on the rate of translation. It may be that cells are capable of exceeding these limits, but that it is not efficient to maintain this capacity.

**Figure 7.**
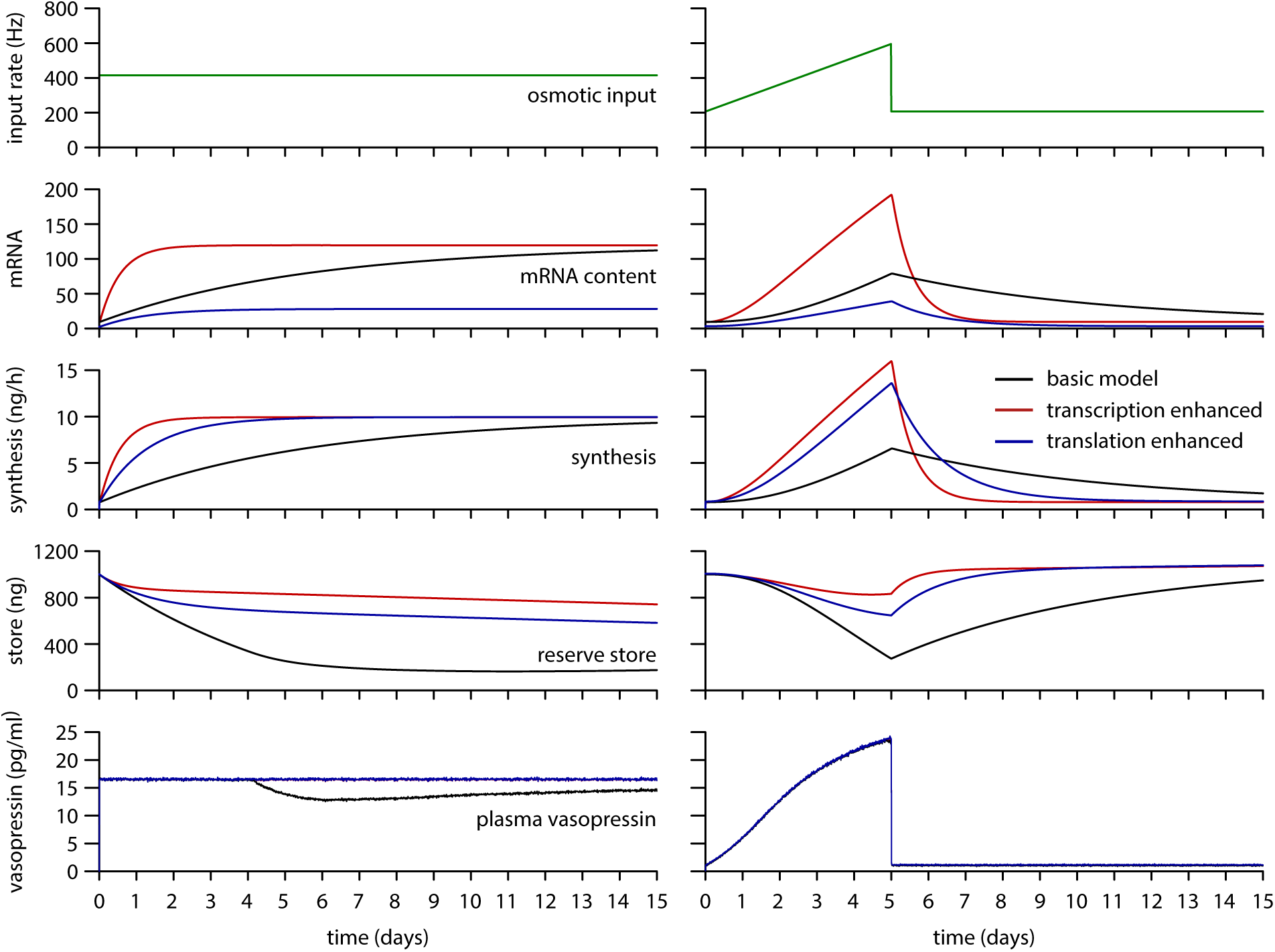
Improving synthesis performance by enhancing transcription or translation. Two enhancements are compared to the basic (black) experimental data fitted model (Figure 5 and 6), attempting to predict what limits the synthesis response *in vivo*. Accelerating transcription (red) produces a much faster upregulation of mRNA content and synthesis rate, reducing store depletion but depending on a larger mRNA capacity. Increasing the proportional translation rate (blue) similarly produces a faster upregulation of synthesis, with a more rapid depletion of mRNA content resulting in a lower equilibrium level. This reduces store depletion but likely depends on a translation rate which is beyond the capability of the cells.

## Discussion

This study is part of a project aimed at understanding how vasopressin neurons function as part of a physiological system on very long time scales. On the surface they appear to perform a very simple signal processing task, producing a plasma hormone signal that linearly encodes osmotic pressure. However, they have many complex features, in particular their distinctive phasic firing, its relationship with the highly non-linear properties of their secretory terminals, and their highly heterogeneous activity levels. The phasic firing is asynchronous and not reflected in the functional signal of their plasma summed secretory output. It appears that the complexity is not about computation, but about being robust, adaptable, and efficient, and maintaining function over lifelong periods of time. This relationship between complexity and function is likely to be true of many neuro-physiological systems, and the experimentally accessible and well-studied vasopressin neurons therefore serve as a very good model system.

Essential to understanding the long term function of neuroendocrine neurons (and other endocrine cells) is the dynamics of hormone storage and synthesis and the focus here has been building and testing a synthesis model to integrate with our existing spiking and secretion model. The new model is built on the work of Fitzsimmons et al (Fitzsimmons *et al*., 1992) which showed that activity-dependent upregulation of mRNA content could best explain the experimentally observed dynamics of store depletion and recovery. The challenge here was to integrate and adapt the model to function without any direct tie between the rates of synthesis and secretion, and for it to function within individual neuron models as part of a heterogeneous population. This has been successful, providing further evidence that the accumulation of mRNA is key to synthesis dynamics. In normal and hypoosmotic conditions mRNA content functions to measure and service current demand. In hyperosmotic conditions it serves as a memory of sustained challenge and following the challenge provides a resource to recover depleted hormone stores.

The mechanisms of the robust new model components presented here are very simple. The key to this was the strongly quantitative properties of the secretion and plasma model. The existing vasopressin neuron model was also further developed here by integrating a new model of hormone diffusion and clearance in plasma, and by refining the quantitative scaling of the existing secretion model, based on previous work in oxytocin neurons (Maícas-Royo, Leng and MacGregor, 2018). The importance of this was in relating rates of secretion to experimentally observed plasma concentrations, thereby accurately simulating hormone store depletion (and recovery) and constraining synthesis rate demands. Initial attempts at building the synthesis model, before the secretion rate scaling had been corrected, and the apparent synthesis rate demands were much higher, used an additional activity-dependent component for the translation rate, shown ghosted in Figure 8. This was not robust, being very sensitive to the balance between parameter values driving the activity dependent transcription and translation components. The version presented here uses only a fixed translation rate, proportional to the mRNA content. Thus, making the model more quantitatively accurate actually reduced the necessary complexity.

**Figure 8.**
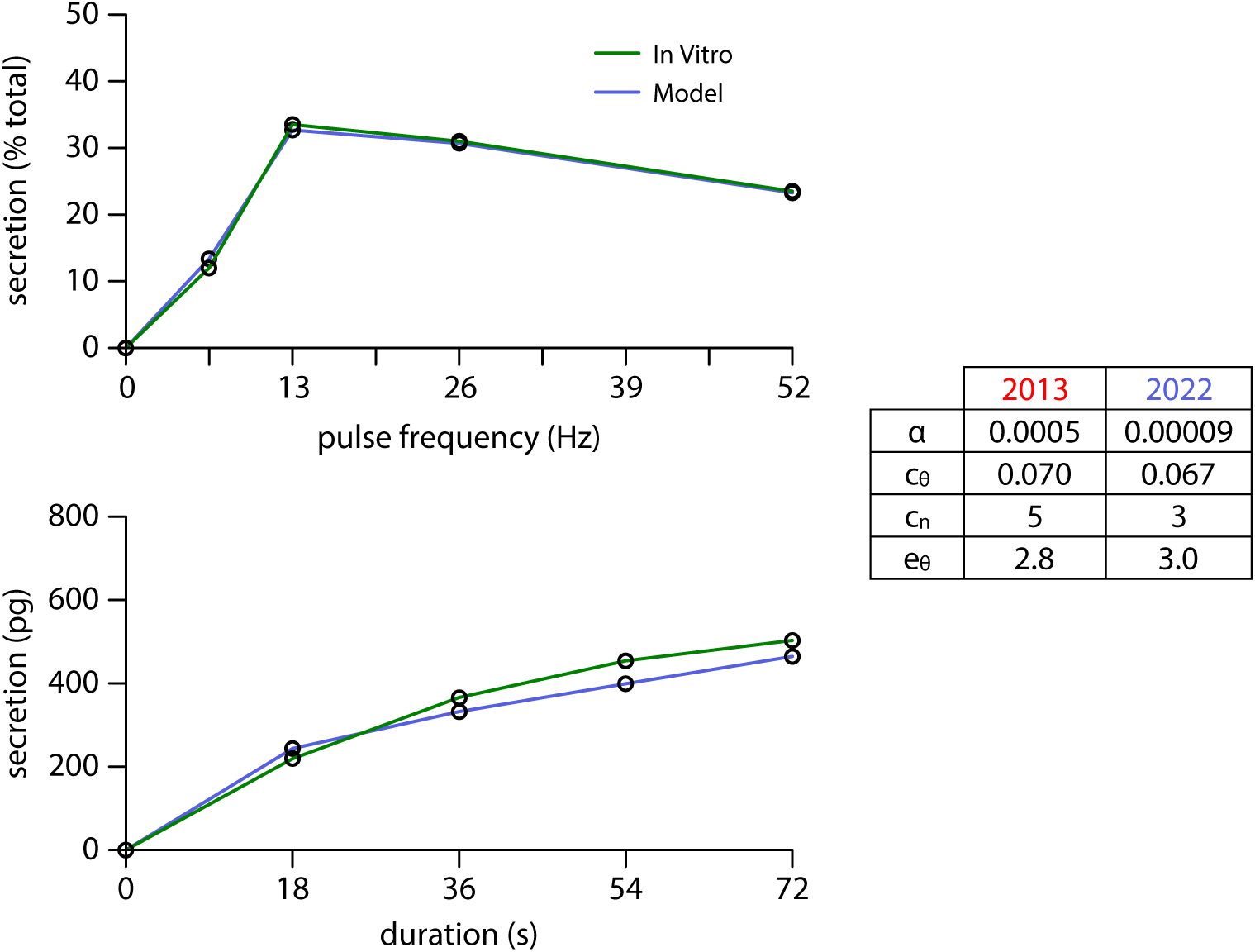
Refinement of the secretion model. The quantitative fit of the published secretion model (MacGregor and Leng, 2013) was improved using more detailed *in vitro* data and better parameter tuning methods based on recent work applying the model to oxytocin neurons (Maícas-Royo, Leng and MacGregor, 2018). The model is fitted to data measuring both frequency facilitation (top panel) and fatigue (lower panel), simulating the in vitro experimental protocols. The changed parameters are given in the table. The major adjustment was to reduce α which scales the rate of secretion to units of pg.

The model also uses a simple representation of the relationship between osmotic stimulus and transcription, making use of the spiking model’s Ca^2+^ variable. Rather than the mechanism necessarily being Ca^2+^-dependent, the necessary assumption here is that transcription closely tracks spike activity. This helps the model to maintain a tracking between the synthesis and secretion rates without any cross-communication. If transcription was more directly driven by synaptic input then the complexities of phasic firing would disrupt this tracking.

Where the model’s behaviour becomes more complex is in the dynamics of stores in a heterogeneous population. Heterogeneous activity levels would be expected to present a big challenge to maintaining the tracking between synthesis and secretion rates and it was surprising how robust the heterogeneous population proved to be. This is partly because heterogeneity as well as adding complexity, also removes some by linearising the relationship between the input and output signals. However, there is some variation across the heterogenous population in how well stores are maintained, suggesting that a statically heterogeneous population will gradually diverge in store content. The simple model tested here puts no limits on the mRNA content or vasopressin stores in individual neurons. These limits are likely to exist in some form and would act to reduce the divergence between neurons, but it does nevertheless seem likely that a static heterogeneous population would struggle to maintain function over long periods. Thus the model here has developed a tool to further examine rather than solve the problem of store divergence identified in the previous work (MacGregor and Leng, 2013).

The alternative is dynamic or regulated heterogeneity. Here we refer to our neurons as a population, connected only at their functional input and output signals. However vasopressin neurons have the ability to communicate through dendritic release of various signals including vasopressin, and potentially act as a network. There is evidence that these signals act to modulate the activity of neighbouring neurons (Gouzènes *et al*., 1998) and it has been proposed that the network might act to cycle activity levels (Scott and Brown, 2010), letting rested neurons replace those that have been more active and become depleted. Such a mechanism would also contribute to the lifetime robustness of the system by compensating for lost neurons. The question for this that arises from the work here is what measure would regulate the dendritic signals? For the same reasons that synthesis rates are not thought to be directly coupled to secretion (distant and distributed stores), it would be difficult to directly measure store depletion. Do the stores available for dendritic release deplete sufficiently in parallel to the peripheral stores? Could elements of the synthesis mechanism also regulate dendritic signalling?

One of the main limitations in interpreting the results here is the highly simplified simulation of the prolonged osmotic challenge. The linear ramp is based on experimental data measuring vasopressin concentrations, osmotic pressure, and/or plasma Na^+^ which show mostly linear increases with time during an osmotic challenge over at least three days. After three days however, these increases tend to slow, probably as the high sustained vasopressin output, and regulation of other elements involved in osmotic homeostasis, such as salt excretion, achieve some sort of new equilibrium. There is also evidence of this in the data measuring store content, where the rate of depletion appears to fall towards the latter part of the challenge. We began addressing this here in the model using a non-linear input ramp, but a much better approach in terms of gaining understanding would be to integrate the neural population model into a simple system model of osmotic homeostasis, providing feedback between the vasopressin output and the osmotic input signal.

Another assumption here is the simple linear encoding between osmotic stimulus and the rate of synaptic input. Recent work in oxytocin neurons (Maícas Royo, Leng and MacGregor, 2019) modelling osmotic stimulus in more detail, to simulate experiments in which plasma oxytocin was measured in response to Na^+^injections or infusions, supports this. The linear encoding assumption was sufficient to closely match experimental plasma concentrations with the model, and it is reasonable to assume similar in vasopressin neurons. The exceptions to this are likely to be in special conditions such as pregnancy.

The work here has modelled the activity-dependent hormone synthesis mechanisms of vasopressin neurons and integrated this into a model of spiking and secretion, further refined and developed to accurately simulate plasma vasopressin concentrations in response to dynamic osmotic stimuli. It has shown that the idea of synthesis driven by the regulation of mRNA content remains robust without any assumption of synthesis directly coupled to secretion, and within the complexities of population heterogeneity. The model provides a strong base for future work exploring the mechanisms that coordinate vasopressin neurons as a network to maintain function over lifelong periods of time, including investigation of the dysfunction of these systems.

## Methods

### The Spiking Model

Many vasopressin neurons when stimulated generate a distinctive phasic pattern of spiking, alternating between sustained bursts and silences lasting tens of seconds. This is modelled here using an integrate-and-fire based model (MacGregor and Leng, 2012) modified to include a set of activity-dependent potentials that modulate excitability to shape spike patterning and generate an emergent bistability, matching the observed phasic firing and detailed spike patterning measured using analysis such as the inter-spike interval (ISI) histogram and hazard function (Sabatier *et al*., 2004). Importantly the model also matches the changes in the phasic spiking that occur in response to a changing input signal.

The excitability modulating potentials include a hyperpolarising afterpotential (HAP), a fast depolarising afterpotential (DAP), and a slow after hyperpolarisation (AHP). Each of these is modelled using a single variable that is step incremented with each spike and decays exponentially. This simple form has proven sufficient to produce close quantitative matches to experimentally measured spike patterning and is used here for the activity dependent elements of the model.

The phasic firing mechanism uses a slow DAP based on a Ca^2+^ inactivated hyperpolarising K^+^ leak current *V*_L_ (i.e. an activity-dependent depolarisation generated by switching off a hyperpolarisation). This is modulated by two opposing step-and-decay variables representing spike generated Ca^2+^ entry, and dendritic dynorphin release, which slowly accumulates to oppose the action of Ca^2+^ and reactivate the K^+^ leak current, eventually terminating a burst and sustaining the period of inter-burst silence.

With more detail in (MacGregor and Leng, 2012; Maícas Royo *et al*., 2016), the spiking model is summarised by:

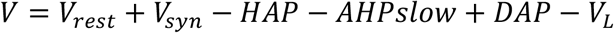

where*V* is the membrane potential, *V*rest is the resting potential, and *V*syn is the summed synaptic input signal described below. AHPslow is the renamed AHP of (MacGregor and Leng, 2012) to distinguish it from the medium AHP of (Maícas Royo *et al*., 2016). The default parameters are given in Table 1.

### Synaptic Input Signal

The osmotic stimulus is encoded by a mixed train of excitatory and inhibitory post-synaptic potentials (EPSPs and IPSPs). Mixed synaptic input contributes to producing a more linear spiking response to an increasing input signal (Leng *et al*., 2001; Maícas Royo *et al*., 2016). This synaptic input signal *V*_syn_ is simulated using a Poisson random process to generate small (3 mV) exponentially decaying positive and negative perturbations to the membrane potential. The proportion of inhibitory to excitatory PSPs uses a fixed value of 0.75, reduced from the previous 1.0 ratio, based on more detailed modelling of magnocellular neurons (Leng, Leng and MacGregor, 2017). The strength of the stimulus is represented by the mean EPSP rate *I_re_*.

### Secretion and Plasma Model

Good quantitative scaling is essential to understanding the qualitative properties of the neurons, and relating mechanism to function. A necessary element for understanding the synthesis mechanisms is to be able to relate the input signal and spiking activity to the output plasma concentration, in order to constrain the rates of secretion and synthesis. Plasma concentration is the most accessible measure of secretion in the experimental data. Our previous work developing the vasopressin secretion model used a simple, single volume estimate of the relation between secretion rate and plasma concentration (MacGregor and Leng, 2013). More recently we adapted the secretion model to oxytocin neurons and integrated a new model of plasma diffusion and clearance (Maícas-Royo, Leng and MacGregor, 2018) which is able to accurately predict experimental measurements of oxytocin plasma concentration in response to both an acute stimulus (CCK injection, (Maícas-Royo, Leng and MacGregor, 2018)) and slower osmotic challenges (Maícas Royo, Leng and MacGregor, 2019).

The plasma model’s volume, clearance, and diffusion rate parameters were fitted using experimental data testing exogenous infusions of oxytocin (Ginsburg and Smith, 1959; Fabian *et al*., 1969). It models peripherally secreted oxytocin as distributed between plasma and extravascular fluid (EVF) compartments, with roughly similar volumes (8.5 ml and 9.75 ml respectively for a 250g rat), diffusing between the two according to the concentration gradient with a time constant estimated by the experimental data. Clearance is a single component from the plasma, representing the total clearance from the kidneys and other organs. There are no equivalently detailed data available for vasopressin, however the vasopressin peptide has a very similar size and transport properties, and we assume that the same volumes and diffusion can be applied to vasopressin.

There are differences however in the clearance rates, mainly due to the added component of bound vasopressin cleared at the kidneys. Experiments measuring the stable plasma concentrations in response to sustained infusions of oxytocin and vasopressin (Robinson *et al*., 1989) estimated the total clearance rate of vasopressin as almost exactly double that of oxytocin. This is consistent with previous experiments that show higher oxytocin concentrations in response to the same stimulus (Dogterom, Van Wimersma Greidanus and Swabb, 1977; Windle *et al*., 1993). Thus we modified the plasma model by reducing the clearance half-life parameter from 68s to 34s. Combined with the diffusion component this produces an overall clearance half-life of ~51s. This matches the estimate of (Ginsburg and Heller, 1953) but is shorter than other estimates of 120s (Czaczkes and Kleeman, 1964).

As well as adding the plasma model we also refitted the existing vasopressin secretion model (MacGregor and Leng, 2013) using the same technique and equivalent data as used to fit the oxytocin secretion model (Maícas-Royo, Leng and MacGregor, 2018). The improved fits (Figure 8) reduced the scaling of secretion per spike (parameter α) by a factor of seven, consistent with the smaller total functional volume estimate of the improved plasma model (18 ml reduced from 100ml).

### The Synthesis Model

The development of the synthesis model tested many more complex forms than those presented here. Our aim was to produce a concise and robust model which is sufficient to make a close qualitative and quantitative match to the available experimental data. The new model adds only two new variables to the integrated vasopressin neuron spiking, secretion, and plasma model, representing the rate of transcription, and the mRNA store (Figure 9).

**Figure 9.**
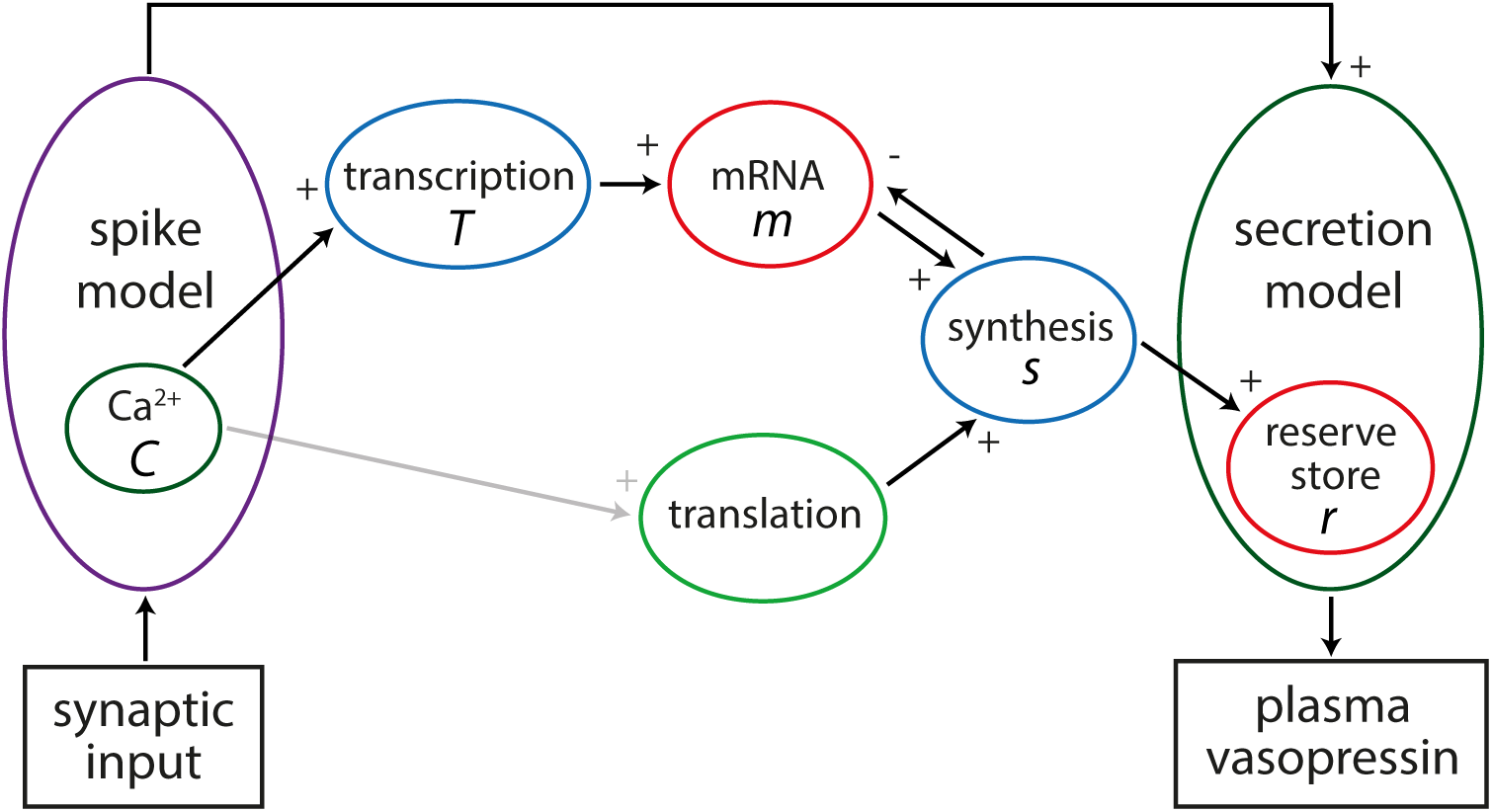
The integrated synthesis model. The spiking model, stimulated by synaptic input that encodes osmotic pressure, drives the synthesis model through its Ca^2+^ variable *C*. This regulates the transcription rate *T* which increases the store of vasopressin mRNA *m*. The synthesis rate *s* is proportional to *m* using a fixed translation rate, and also depletes *m*. The ghosted link showing activity-dependent regulation of translation was not used in the results here. The synthesis model is coupled to the secretion model through the charging of its reserve store *r*.

Transcription is upregulated with the osmotic stimulus. Without any quantitative data available we have not attempted to make any detailed model of the proposed cAMP or other messenger dependent pathway. The essential property of this mechanism is that it needs a sustained activity-dependent signal, acting on a much slower timescale than the rapidly changing electrical activity of the neuron. Informed by previous experience of signal transduction across timescales in modelling circadian and circannual rhythms (Macgregor and Lincoln, 2008) this uses a two-step process. The spiking model’s existing activity-dependent Ca^2+^ variable (*C*) is used, relative to basal Ca^2+^ (*C_rest_* = 113 nM), to drive the transcription rate (*T*), which increases in proportion to *C* at rate 0.001 *k_T_* units per s, and decays exponentially with half-life λ*T* = 1000s:

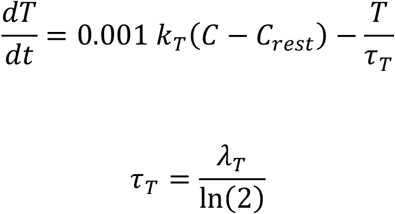

The half-life is very approximate, chosen to produce a slowly changing value that reaches an equilibrium proportional to the osmotic input stimulus (Figure 2). The 0.001 scaling factor produces a more convenient scale for parameter *k_T_*.

The mRNA store (*m*) accumulates at a rate proportional to *T*. Contrary to the Pittsburgh model, it has no explicit exponential decay component, but is depleted in proportion to the synthesis rate (*s*). However, *m* does decay exponentially when translation is at a fixed rate proportional to *m* (parameter *tl_basal_*), as it is in the results here.

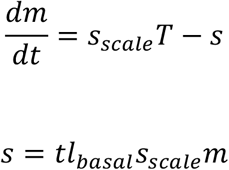

The model includes functional timescales ranging from ms to days, and parameter *S_scale_* is used to scale the rates between the spiking and secretion model and the synthesis model. The results here use a fixed value, but it was convenient for testing to be able to use this parameter to accelerate the synthesis timescale.

The reserve store *r* equation from the secretion model (eqn 8 in (MacGregor and Leng, 2013)) was modified to add the synthesis component:

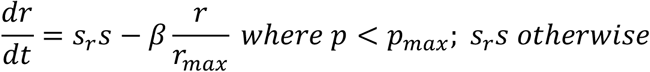

where parameter *S_r_* scales the synthesis units to pg units of stored vasopressin, and parameter β is the refill rate of the secretion model’s releasable pool*p*. The default parameters are given in Table 2. To simulate a transport delay between synthesis and the store, *s* in this equation was replaced *S_delay_*, using the recorded value for *s* from an earlier timestep.

**Table 2:** Default Synthesis Model Parameters

| Name | Description | Value (units) |
| --- | --- | --- |
| $k_T$ | transcription upregulation rate | 0.33 ($T$ units $s^{-1}$) |
| $\lambda_T$ | transcription rate half-life | 1000 (s) |
| $tl_{basal}$ | fixed translation rate | 0.7 |
| $s_{scale}$ | synthesis time scaling factor | 0.000003 |
| $s_r$ | vasopressin scaling factor | 1.1 (pg per $s$ unit) |

### Population Simulation and Heterogeneity

The population was simulated by running in parallel 100 copies of the coupled spiking, secretion, and synthesis model, with the secretion rate outputs forming a summed input to the single plasma model. A heterogeneous population is generated by randomly varying the proportional input rates (synaptic input density *I*_syn_) for each neuron using a lognormal distribution with mean = 0 and standard deviation = 0.25. This approximates the highly heterogeneous range of spiking rates observed experimentally (MacGregor and Leng, 2013). The stimulus is then represented by the population input rate *I*_pop_ and individual neuron input rates are generated using:

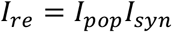

### Implementation

The differential equations were integrated using the first order Euler method. We can do this safely since the step size (1 ms) inherited from the spiking model is much smaller than any of the time constants in the model. Using the same fixed time step makes it simple to couple the synthesis model, and the secretion and plasma models, to the integrate-and-fire based spiking model. The modelling software was developed in C++, using the open source wxWidgets graphical interface library. Each neuron runs as a duplicate integrated spiking, secretion, and synthesis model thread, with secretion rates feeding into a single plasma model thread. A typical run of the full model, simulating 22 days of activity for a population of 100 neurons takes ~18 minutes on an AMD Ryzen 9 5900X 12-core processor.

The model source code, and software, compiled for Windows PC, are available at https://github.com/HypoModel/MagNet/releases.

## Supporting information

Figure S1 osmotic challenge data

osmotic challenge data tables

## Acknowledgement

Professor Gareth Leng is gratefully acknowledged for his contribution to discussions on the project and advice on editing the manuscript.

