## Supplementary figures and images for "Modelling vasopressin synthesis and storage dynamics during prolonged osmotic challenge and recovery based on activity dependent upregulation of mRNA transcription"

### Figure S1 osmotic challenge data

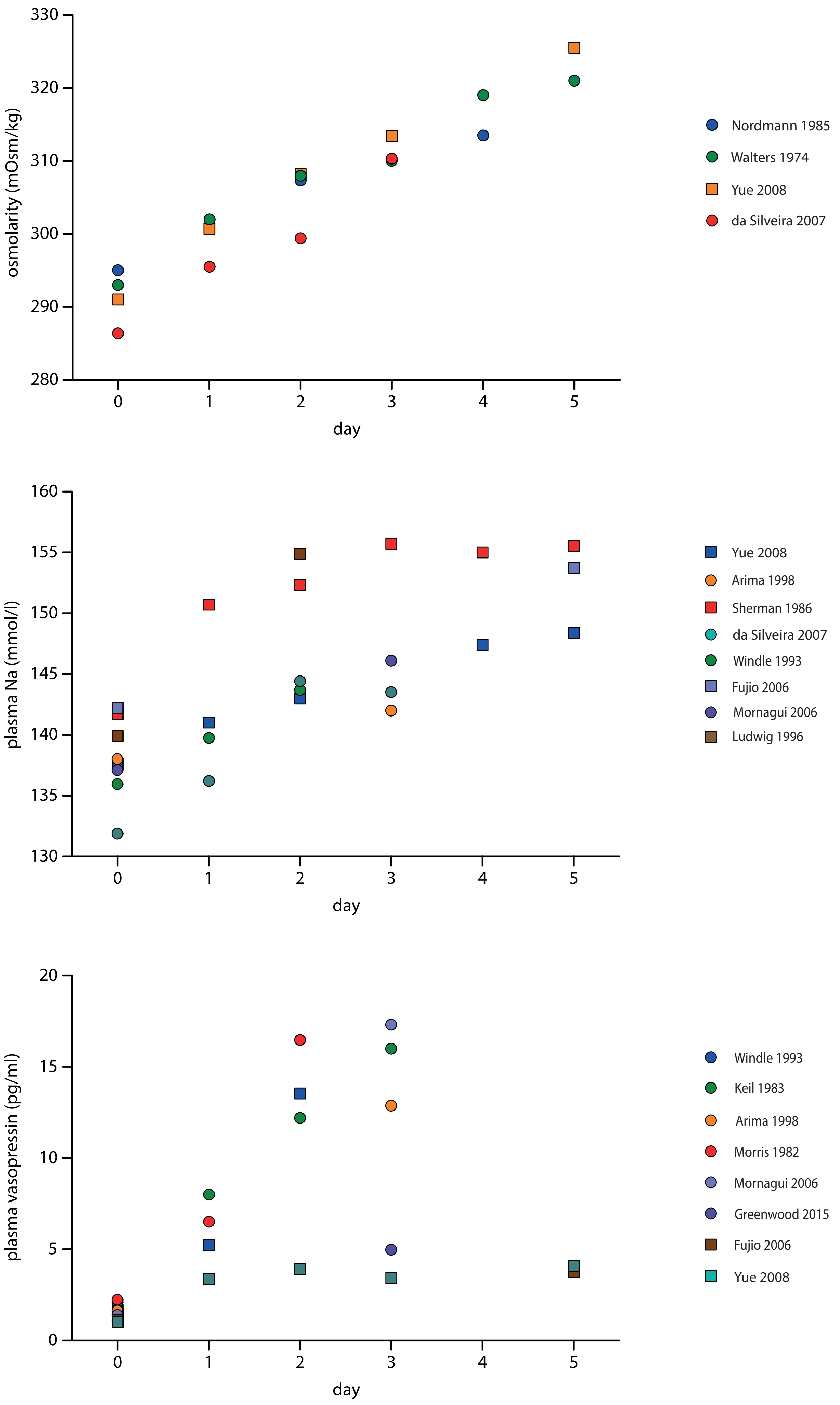

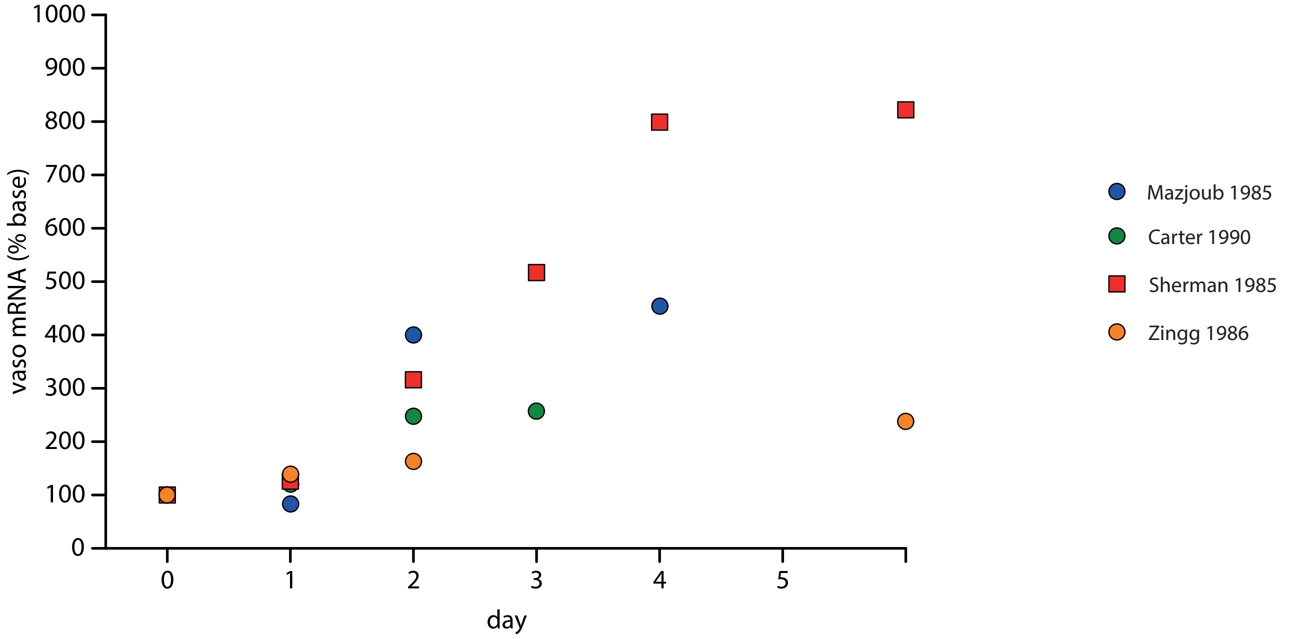

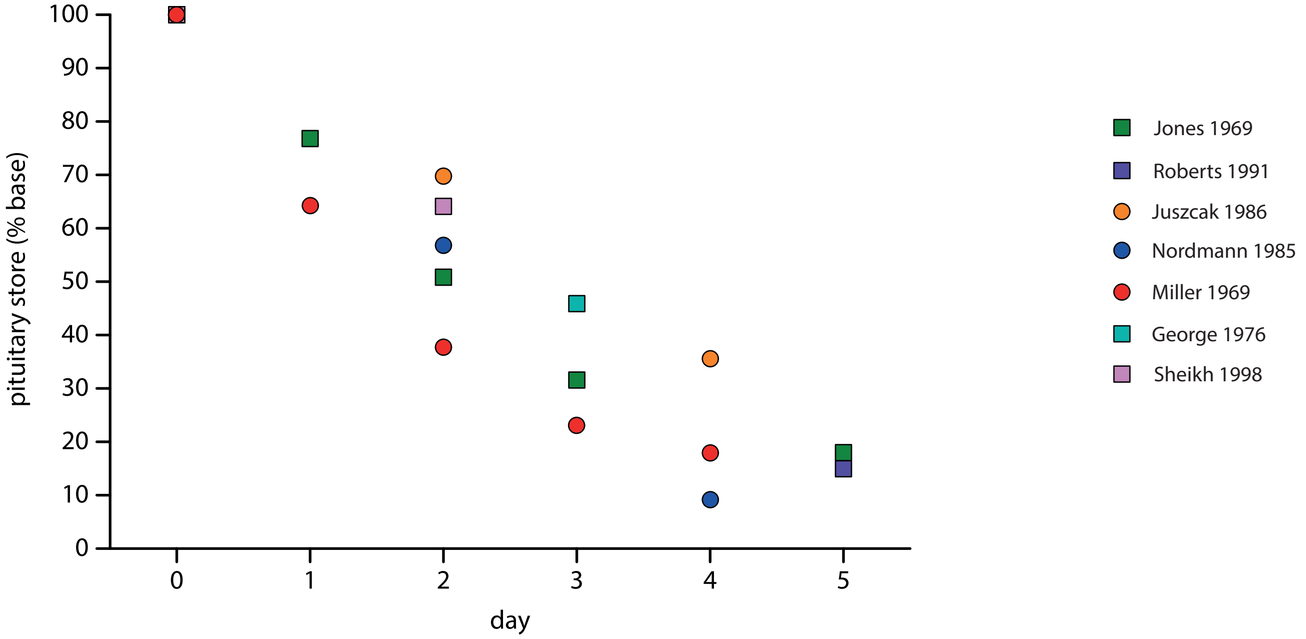

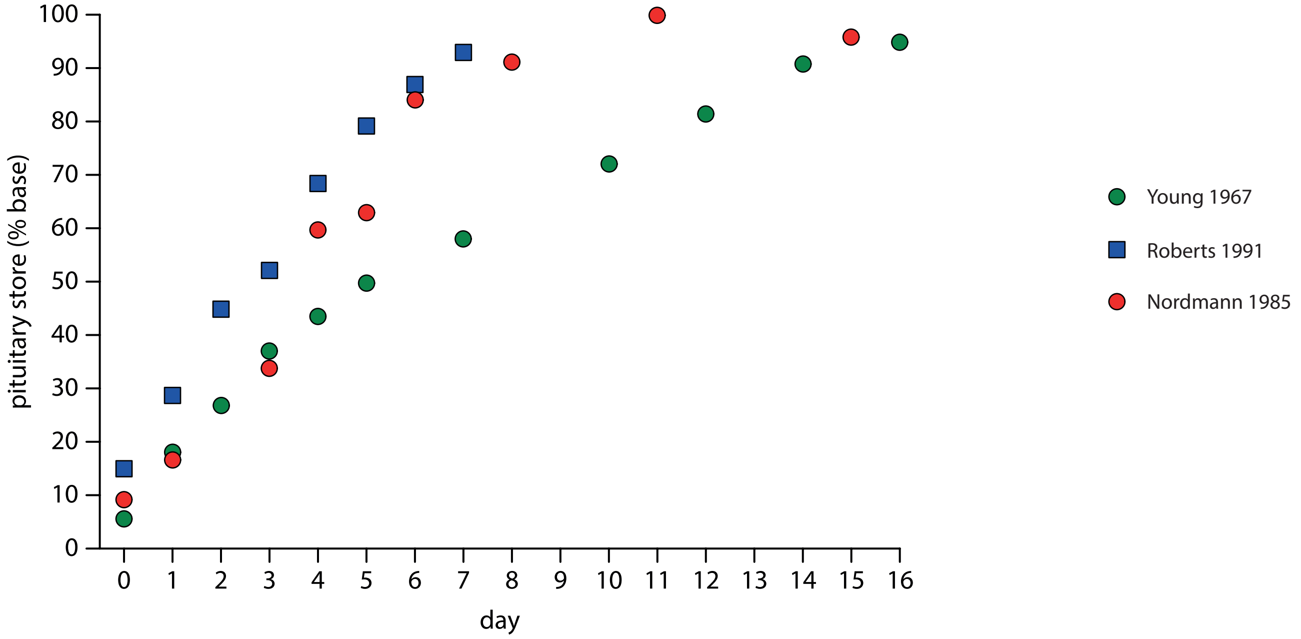
